# Biplanar nulling coil system for OPM-MEG using printed circuit boards

**DOI:** 10.1101/2025.02.19.638883

**Authors:** Mainak Jas, John Kamataris, Teppei Matsubara, Chunling Dong, Gabriel Motta, Abbas Sohrabpour, Seppo P. Ahlfors, Matti Hämäläinen, Yoshio Okada, Padmavathi Sundaram

## Abstract

Optically pumped magnetometers (OPMs) are a promising sensor technology for non-invasive measurement of human electrophysiological signals, in particular the magnetoencephalogram (MEG). OPMs do not need cryogenic cooling and can be placed conformal to the subject’s scalp, thus greatly reducing the sensor-to-source distance and improving signal sensitivity. OPMs, however, require near-zero background magnetic field to achieve linearity and minimize signal distortion. Prior work has proposed the use of biplanar field nulling coils to remove the uniform and gradient components of the background magnetic field. Biplanar coils have been expensive to construct, involving tedious error-prone manual winding of over 1000 m of copper wire. In this work, we designed and fabricated background field nulling coils (three uniform and three gradient components) on two-layer Printed Circuit Boards (PCBs). We used an open-source software (`bfieldtools`) to determine the current loops needed to produce the target magnetic field in a 50-cm-diameter spherical volume. We developed a software-based approach to connect the discrete current loops into a continuous conducting path traversing the two layers of the PCB. For ease of manufacture, the designed (1.5 × 1.5 m^2^) coils were cut along the symmetry axis and printed as pairs of 1.5 × 0.75 m^2^ PCBs (2 oz Cu), soldered together and mounted on a sliding aluminum frame. The efficiency of the coils (1.3 - 7.1 nT/mA) was similar or higher than previously reported in the literature. We mapped the field inside the target region after field nulling inside our single-layer shielded room and were able to reduce the largest component of the background field from 21 to 2 nT. Using our nulling coil system, we were able to operate OPMs in a lightly shielded room (background field varying from 6.5 to 108 nT in the floor-to-ceiling direction) to record somatosensory evoked fields (SEFs) comparable to those measured using SQUID-based MEG in a 3-layer shielded room. We disseminate the software and hardware as an open-source package `opmcoils`. This work will facilitate access to more affordable field nulling coils for OPM-MEG and help to realize the potential of OPM-MEG as an accessible sensor technology for use in human neuroscience.

## 1. Introduction

Optically Pumped Magnetometers (OPMs) are a promising new sensor technology for magnetoencephalography (MEG). They allow the measurement of weak magnetic fields from the brain at room temperature (Boto et al., 2018). Unlike conventional cryogenic Superconducting Quantum Interference Device (SQUID) sensors, OPMs do not require a fixed gantry and can be placed in a flexible manner on the subject, thus expanding the potential applications of MEG. OPMs have been used, e.g., in naturalistic paradigms involving movements (Boto et al., 2016) as well as in pediatric MEG studies (Feys et al., 2022). Placed on-scalp, the OPM sensors may also improve the spatial resolution obtainable using MEG (Iivanainen et al., 2021; Jas et al., 2021). Despite the potential of OPMs, they have not been widely adopted (Bagić et al., 2023), partly because OPMs require a near-zero background field for optimal operation which remains a challenging problem (Alem et al., 2023; Borna et al., 2022). The near-zero background field is necessary to avoid saturation and non-linearity of the OPM sensors (Tierney et al., 2019). In addition to cancellation of the uniform field, minimizing the field gradient is important for reducing artifacts due to movement (Brookes et al., 2021).

On-sensor Helmholtz coils built into OPM sensors can null background fields also in closed-loop mode, enabling continuous nulling even when the background field drifts. Even then, the dynamic range of the sensors is limited to ±15 nT in closed-loop mode (Alem et al., 2023). Any sensor movement that changes the background field greater than this dynamic range will cause the sensor to saturate or move outside the linear operating regime (Tierney et al., 2019). On-sensor coils may introduce cross-talk and may be challenging to operate in lightly shielded rooms (Alem et al., 2023; Nardelli et al., 2019).

To address these problems, field nulling coil systems have been developed to minimize the background field (uniform and gradients) inside a target region containing the subject’s head (Holmes et al., 2018, 2019; Iivanainen et al., 2019). Simple Helmholtz coil-based designs (Iivanainen et al., 2019) are efficient, but they are not always appropriate for neuroimaging studies as the coils may obstruct access to the subject. Biplanar “fingerprint” coils have been designed (Boto et al., 2018; Holmes et al., 2018, 2019; Tian et al., 2025) where the current distribution is restricted to a planar surface. They allow field nulling inside a target region using mathematically optimized current distributions (**Fig. 1**). Biplanar coils trade lower efficiency, i.e., magnetic field generated per unit current applied to the coils, for improved usability and access to the subject. Newer systems using matrix coil designs (Holmes et al., 2023) are being developed, but they are less efficient, more complex to operate and more expensive due to the number of independent current drivers required for their operation. Biplanar “fingerprint” coils therefore remain the most common choice for background field nulling in OPM systems.

**Figure 1:**
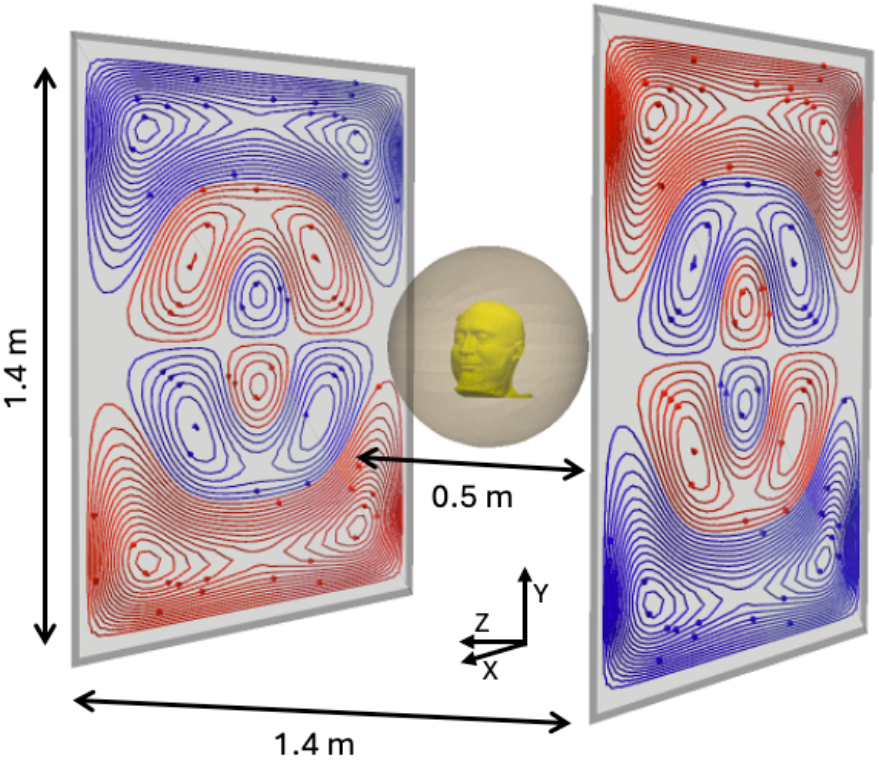
The overall dimensions of our field nulling system. 1.4 × 1.4 m^2^ coils were etched in a PCB pair of size 1.5 × 1.5 m^2^ and separated by 1.4 m. The target region is a sphere of diameter 50 cm.

A challenge with existing biplanar coil designs (Holmes et al., 2018, 2019) is that they involve tedious error-prone manual winding of large quantities (> 1000 m) of copper wire. To construct such a coil, the winding pattern is printed on a sheet of paper, then wire is glued onto it using epoxy, and the assembly is supported using a medium-density fiberboard. To minimize the resistance and thermal noise, a low gauge (thick) wire is used, making winding difficult. To improve ease of construction, enhance reproducibility, and avoid manual winding, our goal was to automate as much of this process as possible. Printed Circuit Boards (PCBs) present a promising solution by allowing high-precision etching of the desired current path. However, PCB-based coils are not straightforward to design with existing tools because they yield disjointed current loops which must be fully connected into a continuous conducting path that traverses the PCB layers.

Here, we present an open-source semi-automated pipeline for manufacturing nulling PCB coils that addresses the practical challenges with existing biplanar coil systems. PCBs allow for precision in manufacturing and easy replication since the design files can be shared. Our open-source package `opmcoils` contains both the Python-based software needed to design the nulling coils, the design files used to print our PCBs, and instructions for installation. We found that the material cost of our system was significantly less than the cost of currently available commercial nulling coil systems. We expect this to contribute to more affordable and easily available field nulling options for OPM-MEG.

In this paper, we outline our process and discuss the challenges we faced in using existing the newly developed tools. We characterized the accuracy and efficiency of the coil system and compared experimental data to theoretical values. We mapped the residual field before and after using the field nulling system. We also demonstrate somatosensory evoked responses recorded with OPMs operating in the environment created by the PCB coils in a one-layer shielded room, providing data comparable to that measured by SQUIDs in a 3-layer room.

## 2. Methods

We developed `opmcoils`, a software tool that provides an application programming interface (API) for coil design and abstracts away the technical implementation details. The purpose was to catalyze future development of the field nulling systems by allowing developers to obtain the current loops, connect them into a continuous path, evaluate the coil efficiency as a function of design parameters (e.g., coil size, inter-coil distance, shielded room), and export the conducting paths to manufacturable PCBs. Below, we describe the fabrication, installation and evaluation of the field nulling PCB coils designed to be used for OPM-MEG in a lightly shielded room.

### 2.1 Pipeline for manufacturing biplanar nulling PCB coils

#### A. Optimizing and discretizing stream functions

The target field method has been widely used in MRI for gradient coil design (Liu, 1998; Martens et al., 1991; Turner, 1986; Yoda, 1990) as well in transcranial magnetic stimulation (Koponen et al., 2017; Sánchez et al., 2018) The underlying physical principles and the optimization problem specific to OPM-MEG nulling coil design are adequately addressed in the literature (Holmes et al., 2018, 2019; Mäkinen et al., 2020; Zetter et al., 2020). We used `bfieldtools`, a Python-based open-source software package for magnetostatic calculations on surfaces of arbitrary shape (Mäkinen et al., 2020; Zetter et al., 2020).

`Bfieldtools` represents magnetic fields using spherical harmonic functions and the unknown current densities using stream functions. The stream functions are restricted to a triangle mesh representing the coil surface. The estimation of the unknown stream function *s* is a convex optimization problem with a quadratic objective function,

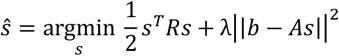

where *R* is the resistance matrix, *A* is the coupling matrix that maps the stream function *s* to the measured magnetic field *b* and *λ* is the regularization parameter. One can think of the minimization of the quadratic term *s*^*T*^*Rs* as minimizing the resistive loss in the coil.

Under the hood, `bfieldtools` uses CVXPY (Diamond & Boyd, 2016) to solve the quadratic optimization problem. Once the estimate ŝ was obtained from `bfieldtools` (e.g. **Fig. 2A**), it was discretized to obtain a certain number of turns (we used N=30, **Fig. 2B**). These optimizations were done for each of the three uniform field coils (*B*_*x*_, *B*_*y*_, *B*_*z*_), and for each of the gradient coils 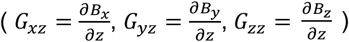. The *z*-axis was defined to be normal to the coil planes, going through the center of the coils, y-axis vertical from floor to ceiling, and x-axis parallel to the coil planes (**Fig. 1**).

**Figure 2:**
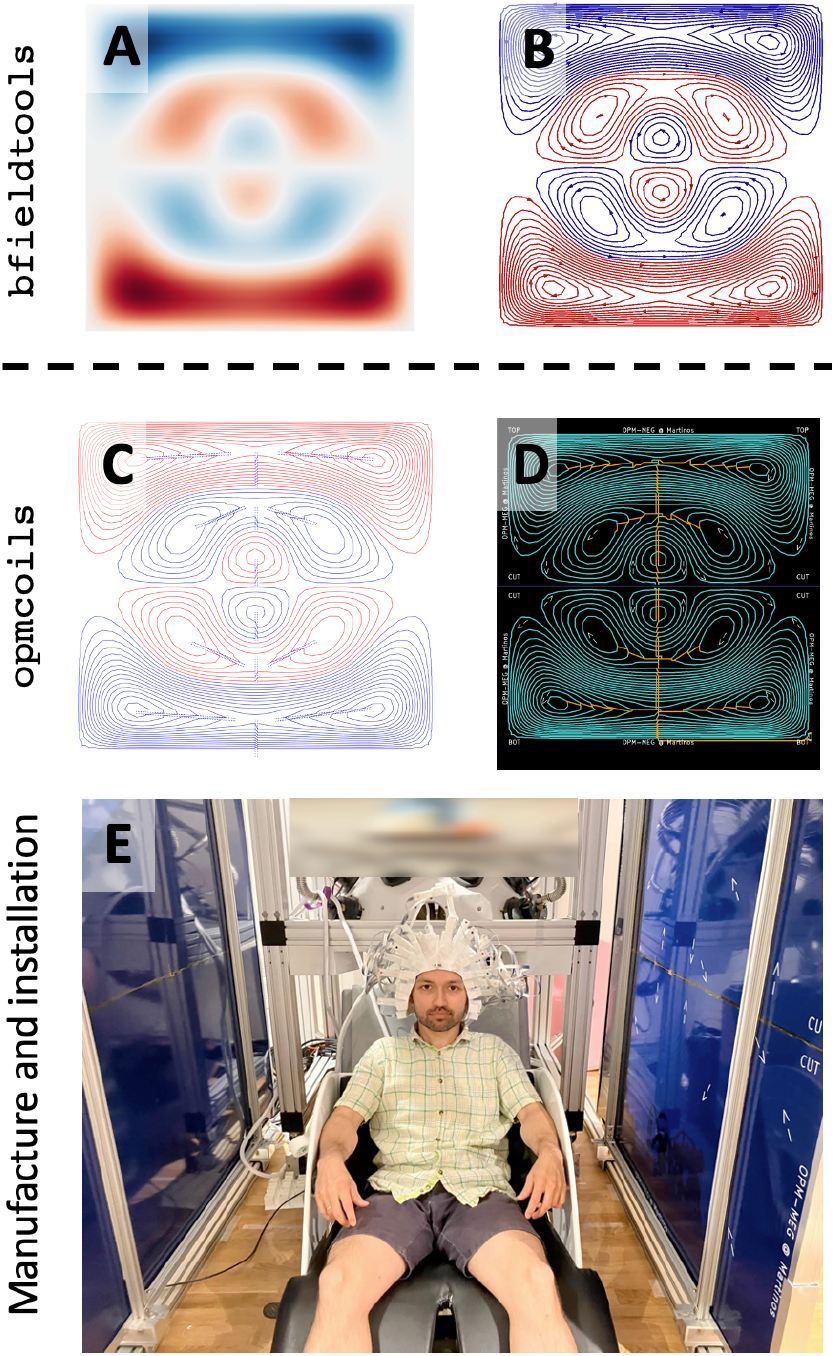
Pipeline for manufacturing biplanar nulling coil system using Printed Circuit Boards. Existing open-source tools only allow optimization and discretization of the stream functions (**A** and **B**) but hardware realization of the optimized loops is expensive and non-trivial. We developed `opmcoils` (**C**-**D**) to enable the development, evaluation and manufacture of biplanar nulling coils. The nulling coils can then be used in OPM-MEG experiments as demonstrated here by the first author (**E**).

We extended the capability of `bfieldtools` to compute the efficiency for the discretized coils with a given number of turns and trace width. Here, coil efficiency was defined as the magnitude of the magnetic field generated per unit current applied to the coil (in units if nT/mA for the uniform field coils or nT/m/mA for the gradient coils). Once the efficiency of the designed coils was found to be sufficient, the remaining challenge was that the discretized current loops were disconnected from each other. In the next section, we describe how these loops were connected into a single continuous path traversing the two layers of the PCB while minimizing stray magnetic fields.

#### B. Connecting current loops

We developed an interactive Python tool to join the discretized current loops into a continuous path. The discretized current loops needed to be either clockwise (**Fig. 2B**: red) or anticlockwise (**Fig. 2B**: blue). Neighboring loops with the same current direction were connected in series along a line segment drawn interactively with our software (**Fig. 2C**). As the current path spiraled inward or outward, it reached the innermost or outermost loop with current in the same direction. Since the additional wire segments used to connect the discrete loops together were not part of the optimized coil design, they were canceled with opposing current paths in the second layer (**Fig. 2D**). At the end of this step, the connected current loops formed “islands” which were disconnected from each other and from the power terminals of the PCB. The final step was performed manually; we exported the connected current loops to KiCad (https://www.kicad.org/), a free open-source electronic design software for PCB manufacturing. We designed the coils for 1.5 × 1.5 m^2^ square PCBs but due to manufacturing size constraints, the coil needed to be printed in two 1.5 × 0.75 m^2^ pieces. Therefore, during export to KiCad, our Python code cut the connected coil traces along the axis of symmetry (vertical cuts for *B*_*x*_, *G*_*xz*_, *B*_*z*_, *G*_*zz*_; horizontal cuts for *B*_*y*_, *G*_*yz*_ coils; **Fig. 3**). Cutting along the symmetry axis meant that at least for the *B*_*x*_, *B*_*y*_, *G*_*xz*_ and *G*_*yz*_ coils, only two solder joints (one for each layer) would be required to connect the two pieces (**Fig. 4C-D, 5C-D**). Only the *B*_*z*_ and *G*_*zz*_ coils needed multiple solder joints to connect the two pieces (**Fig. 4C-D, 5C-D**). Below, we explain the final step in KiCad to create printable design files for PCB manufacture.

**Figure 3:**
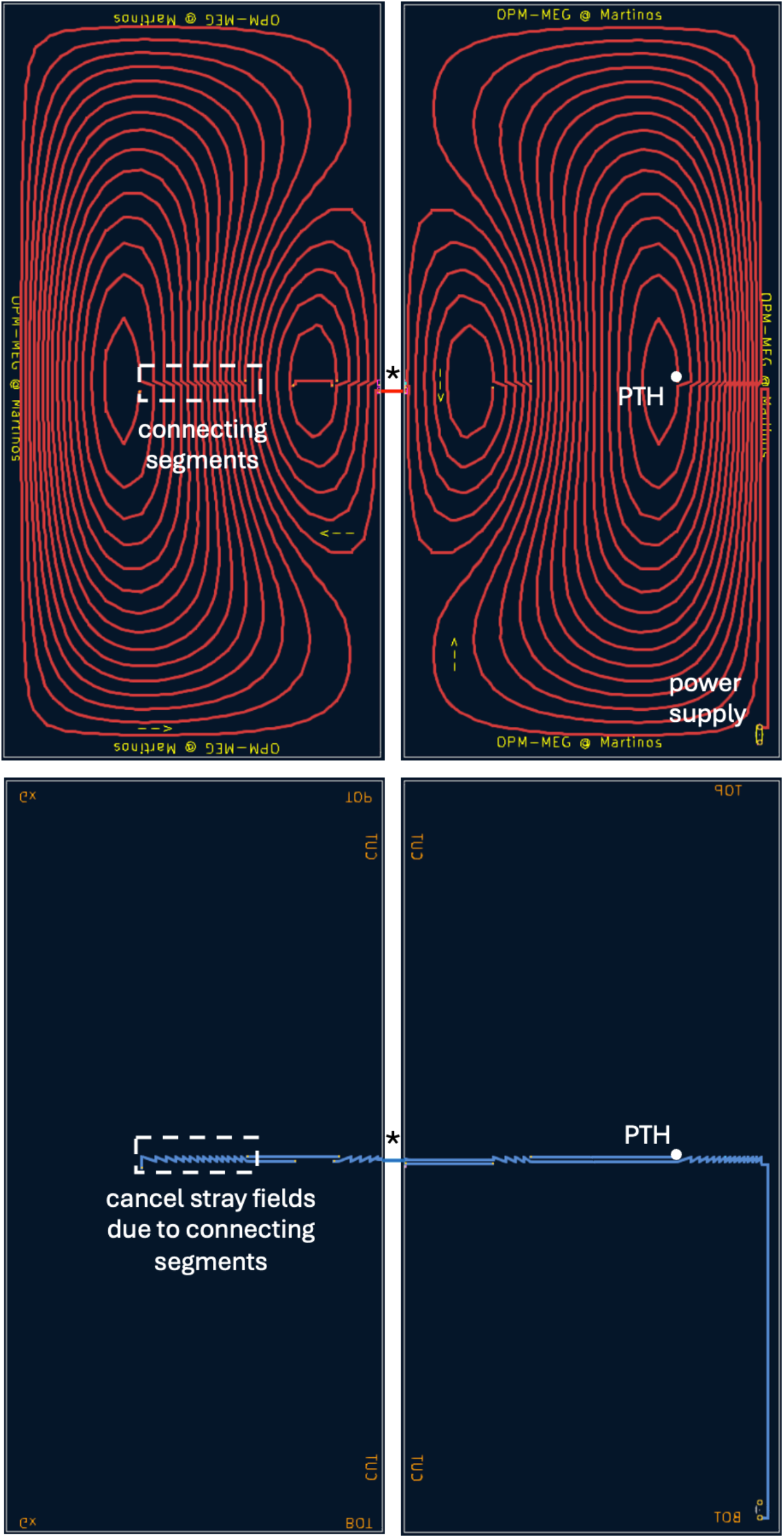
Completed two-layer nulling coil design (*G*_*xz*_) visualized in KiCad, the electronic design software. `opmcoils` generated the wire paths to connect the discretized loops into a continuous conducting path in each PCB. The connecting segments were identical in the front (**top**) and back layer (**bottom**) of the PCB but with opposing current directions, thus self-canceling any additional stray fields. Plated through holes (PTHs) connected the front and back layer at specific locations. The coil was cut along the symmetry axis into two PCBs (left and right columns). During installation, the front and back layers of the PCBs were soldered at designated solder masks (*) to create a complete conducting path.

**Figure 4:**
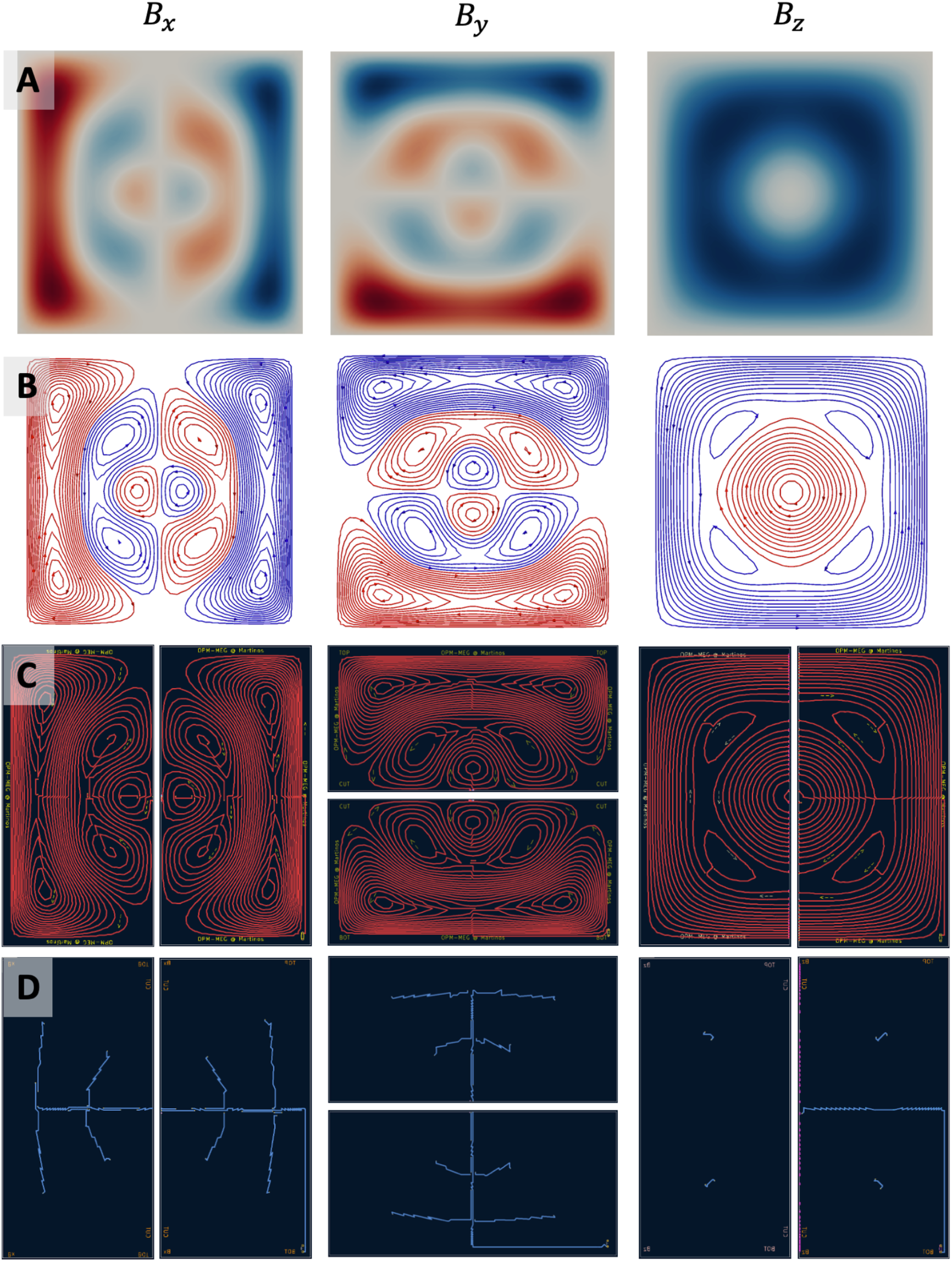
Completed designs for uniform field (B_x_, B_y_, and B_z_) nulling coils starting from optimized stream functions (**A**), discretized current loops (**B**), and connected current paths in the two-layer PCBs (**C** front layer, and **D** back layer)

**Figure 5:**
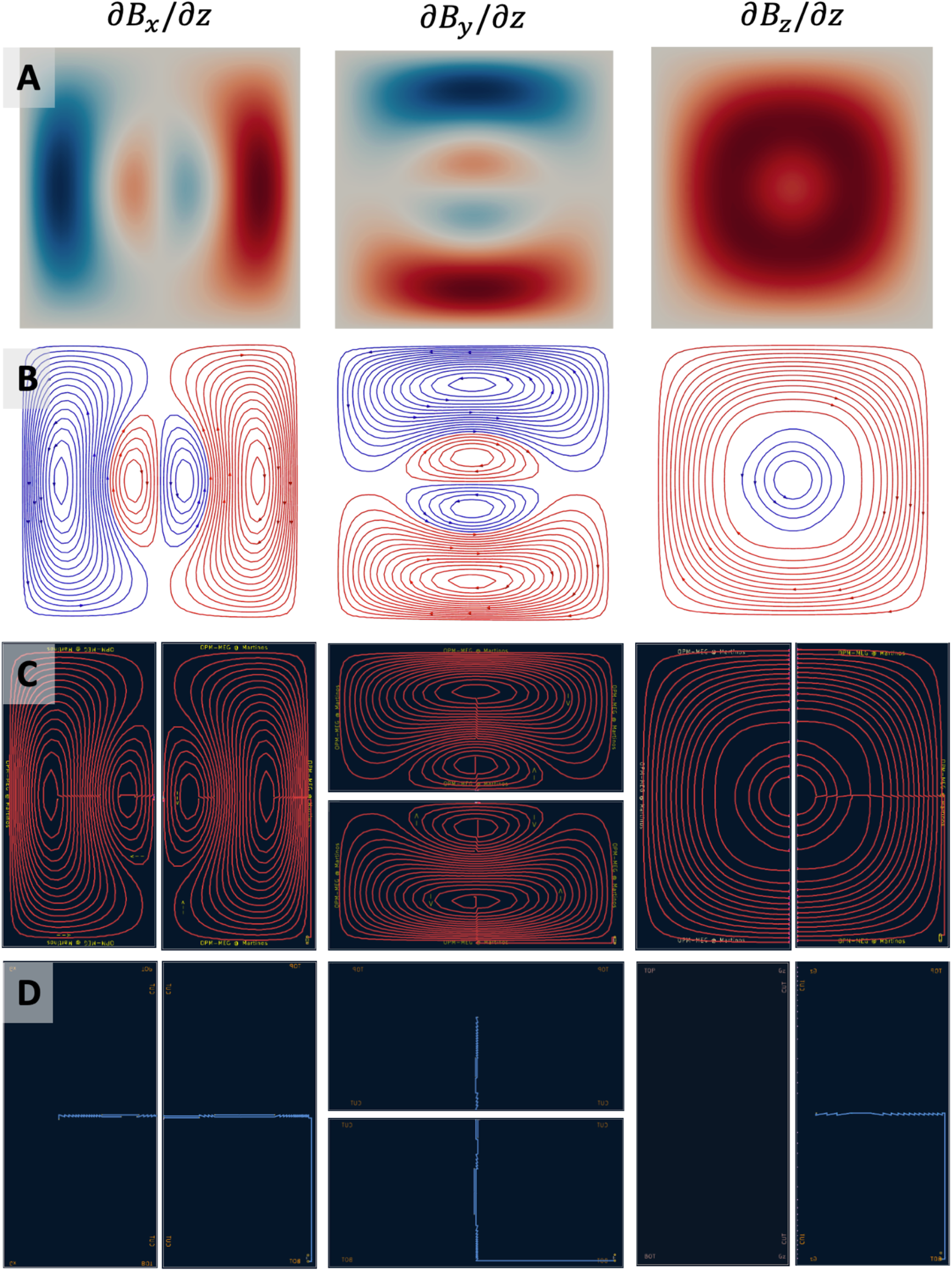
Completed designs for field gradient (*∂B*_*x*_/*∂z, ∂B*_*y*_/*∂z*, and *∂B*_*z*_/*∂z*,) nulling coils starting from optimized stream functions (**A**), discretized current loops (**B**), and connected current paths in the two-layer PCBs (**C** front layer, and **D** back layer)

#### C. Printed circuit board design

We used two-layer PCBs with standard 2 oz copper: each board contained a front copper layer and a back copper layer (**Fig. 3**). The front layer traced the main coil pattern, and the back layer was used to “lift the pen” and move from one set of the current loops to another, without crossing paths on the coil patterns in the plane. We set the PCB trace width to the maximum feasible (5 mm) to minimize the coil resistance. To connect one layer to another, we used 2 mm diameter copper plated through holes (PTHs). The current loop “islands” were connected to each other and to the power terminals on the PCB using the wire drawing tool in KiCad. 5 mm wire segments were drawn along the horizontal or vertical directions with the return paths in the back layer canceling any stray magnetic fields. During the KiCad finishing step, we annotated the coils (text labels indicating cut locations and current directions for the different loops). These annotations were made on the silkscreen layers (one for the front, one for the back) to help with the assembly after manufacture. At the cut edges on front and back, we added solder pads for soldering the boards together during installation. The power terminals were added and aligned across all the coils.

#### D. Manufacturing and installation

The final coil designs were exported to Gerber format files and sent for manufacturing (Shenzhen Hopetime Industry Co., Ltd., Shenzhen, China). Gerber files are a common PCB design format that store the shape and location information for every element in the PCB. Each layer of the PCB (in our case, front and back Cu, front and back silkscreen for annotation and a drill file with the locations of the PTH) was exported into its own Gerber file. A key design consideration was the ability to test and replace coils if necessary. The coils were therefore designed to be stacked together with an 8 mm inter-coil spacing. We accounted for this spacing during the coil design. We first printed and tested the *B*_*y*_ coil pair. Once we were satisfied with the performance of the first coil pair, the remaining coils were printed. For all the printed coils, the two PCB halves were manually soldered together. Since the PCBs were large and heavy, we designed a frame to mount the entire nulling coil system and hold it upright to prevent buckling. Our frame design allows the mounted PCBs to slide on rails depending on the study requirements.

Using the pipeline outlined above, we designed and fabricated six coils: three uniform field coils (*B*_*x*_, *B*_*y*_, *B*_*z*_), and three gradient coils (*G*_*xz*_, *G*_*yz*_, *G*_*zz*_). Since, in a current-free region, ∇ × **B** = 0, our coils can thus also null *G*_*zx*_ and *G*_*zy*_ (Holmes et al. 2018).

### 2.2 Theoretical Estimation of Coil Resistance and Efficiency

We computed the theoretical resistance for the discrete current loops and the joined current path on the PCB for each coil. We used `bfieldtools` to compute the total length of the disconnected discretized current loops and `opmcoils` to compute the total length of the coil which will be printed into a continuous path on the PCB. The resistance of the coil is a function of the total length, the PCB trace width, and the density of copper coating (2 oz = 70 μm). Increasing the copper weighting on a large PCB can be quite expensive but one can easily increase the trace width to the widest feasible (5 mm) without any increase in cost or difficulty in winding. Lower coil resistance reduces thermal noise from the current driver, thus improving the overall quality of the applied nulling field.

The theoretical efficiency (in units of nT/mA or nT/m/mA) of each of the six coils was calculated with and without accounting for the shielded room. We estimated the theoretical efficiency of the coils without the presence of the shielded room. In addition, we used the approach of (Mäkinen et al., 2020; Zetter et al., 2020) to model the effect of field distortions due to the high-permeability mu metal in the floor, walls, and ceiling of our shielded room. We modeled the mu metal as an infinite permeability material that requires an additional boundary condition. The boundary condition was satisfied by setting the scalar magnetic potential at the inner shield to 0 and introducing an equivalent stream function for the magnetic shield:

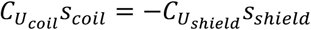

where 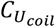 and 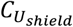 are the magnetic scalar potential coupling matrices. With this equipotential boundary condition, the total magnetic field at the target points was computed as:

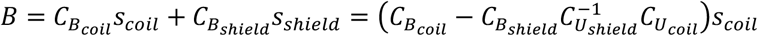

These values were used to calculate the theoretical estimate of the coil efficiency accounting for the effect of the shielded room. Both calculations were incorporated in `opmtools`.

### 2.3 Empirical Measurement of Coil Resistance and Efficiency

We measured the total resistance of each coil pair with a multimeter (Fluke 115 from Fluke, Everett, WA). We used low-noise bipolar constant current drivers (CSB-40, CSB-100 from TwinLeaf, Princeton, NJ) each with three independent outputs. The 100-mA current driver (CSB-100) was used for *B*_*x*_ and *B*_*y*_ coils since they were theoretically less efficient than the other coils. For each coil pair, we measured the background field at the center of the target region as a function of the applied current (0 to 60 mA). The background field was measured using a fluxgate magnetometer (FVM400, Meda, Dulles, VA). The efficiency of the uniform field coils was computed as the slope of the best fitting line to the measurements. Similarly, to measure the gradient, we 3D-printed an array of sensor holders spaced 5 cm apart between -15 cm and 15 cm along the z axis (x=0, y=0; middle of the nulling coils). The efficiency of the gradient field coils was computed by varying the measured gradient as a function of the applied current (0 to 24 mA).

### 2.4 Mapping the remnant field inside the target area

We used OPMs to map the spatial profile of the residual magnetic field in the target zone in the x-z plane after the uniform field coils were applied, sampled at a 5 cm resolution in a 20 cm^2^ region (**Fig. 6**, bottom). Sensor holders for measuring the x, y, and z components of the field were 3d-printed along with a reference pegboard to align the measurements (**Fig. 6**, top). The measured magnetic field was the remnant field nulled by the on-sensor coils in the OPMs (Alem et al., 2023). We used 19 OPMs (Fieldline Gen 2, Boulder, CO) with their sensitive axes along the measured field component. The effect of time-varying drifts in the background field were minimized by measuring the remnant field simultaneously from all the OPM sensors. The bias in the remnant field estimate for each OPM sensor was measured by averaging two observations: one in the original position and another with the sensor flipped along its sensitive axis. The field maps were produced after correcting for this (measurement) bias.

**Figure 6:**
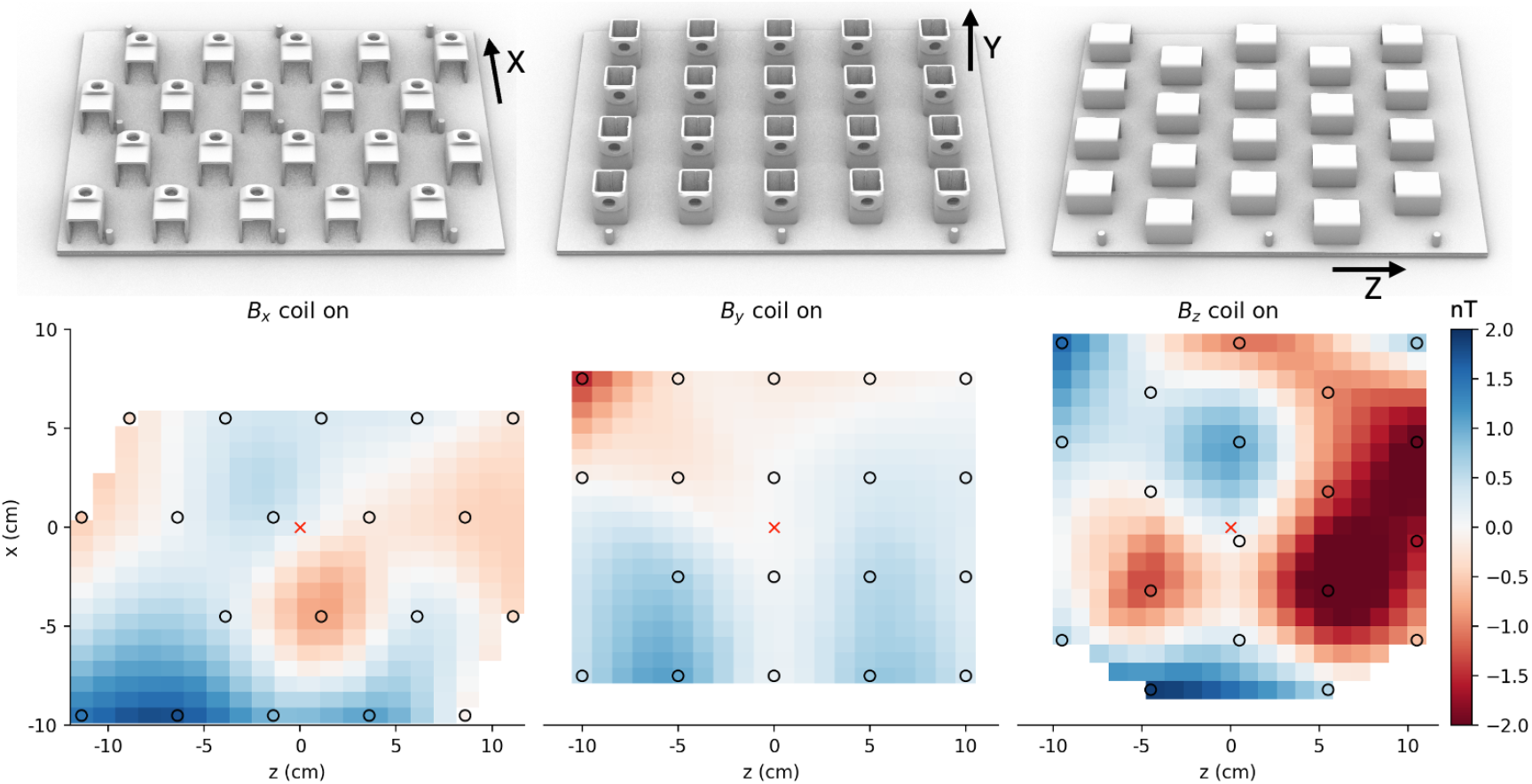
Spatial maps of the residual field in the x, y, and z directions. **Top:** Model of the 3d-printed sensor holders used for mapping. A pegboard was used to align the maps across the three measurements. **Bottom:** Spatial maps after turning on the *B*_*x*_, *B*_*y*_, and *B*_*z*_ nulling coils respectively. The maps were centered at the target region (red cross) in the x-z plane. The positions of the centers of the OPM’s vapor cells are shown in black circles. Field values elsewhere were interpolated.

### 2.5 Estimating optimal applied current on a helmet

An automated procedure was devised to set the currents to minimize the background field in the OPM sensors inside a helmet. Following the method of (Iivanainen et al., 2019), we used a procedure that does not require any additional sensors to determine the applied current. The fields *b* in OPM sensors corresponding to the nulling coil currents *I* were represented using a coupling matrix *M*, such that *b* = *MI*. By driving the nulling coils one at a time with known currents, the columns of coupling matrix *M* can be experimentally determined. Then, to null a measured field *b* with known coupling matrix *M*, the driving currents can be estimated from *I* = *M*^‒1^*b*.

The automatic nulling procedure zeroed the fields at target sensors without explicitly estimating the uniform and gradient components of the background field. This method worked best when the coils produced dissimilar measurements so that the coupling matrix *M* had a reasonably low condition number (Iivanainen et al., 2019). Accordingly, we placed the sensors in a subject-specific helmet (while not being worn by the subject) where the sensors are oriented in multiple directions. The residual field on applying the optimal current was measured using OPMs and mapped on the helmet surface.

### 2.6 MEG measurement and analysis of somatosensory evoked fields (SEF)

We performed a median nerve stimulation experiment (Cruccu et al., 2008) on one healthy adult subject (male, 35 years old). The participant provided written informed consent. The study design, protocol, and consent form were approved by the Massachusetts General Hospital Institutional Review Board. We first performed the experiment using conventional SQUID-based MEG (306-channel Triux neo system, MEGIN Oy, Finland). The subject was asked to stay awake for the duration of the experiment (4 mins) and the median nerve on the right wrist was stimulated with inter-stimulus interval of 500 ms. We repeated the experiment in OPM-MEG (19 sensors, FieldLine Gen 2, FieldLine, Boulder, CO) using the same paradigm. The field nulling coils were operated, and the currents were set using the approach in **Section 2.5** (as demonstrated in **Fig. 2E**). A subject-specific helmet was created by expanding the subject’s FreeSurfer reconstructed scalp surface by 2 mm. For co-registration, we performed digitization using a Fastrak Polhemus system (Polhemus Corp., Colchester, VT). The head-to-MRI transform was estimated by aligning 3 digitized fiducial points (nasion, left and right pre-auricular points) with the same landmarks identified from the MRI. A device-to-head transformation was estimated by aligning 5 pre-determined reference points on the subject-specific helmet with the corresponding points in the 3D model of the helmet. 16 sensors were placed in the helmet holders covering the contralateral somatosensory cortex.

The data was high-pass filtered at 4 Hz and low-pass filtered at 150 Hz to preserve the shape of early N20 components. Due to inherent noise in the OPM sensors, a low pass filter was necessary. In SQUID-MEG, background noise was removed using the signal space projection (SSP) method (Uusitalo & Ilmoniemi, 1997). All analysis was performed using MNE-Python (Gramfort et al., 2013).

## 3. Results

### 3.1 Coil Designs

**Figs. 4** and **5** show the stream functions, discretized current loops, the connected current paths from `opmcoils`, and the front and back copper layers of the two PCB halves in KiCad for each of the six coils (*B*_*x*_, *B*_*y*_, *B*_*z*_, *G*_*xz*_, *G*_*yz*_, *G*_*zz*_). We note that the *G*_*xz*_, *G*_*yz*_, *G*_*zz*_ coils were similar to *B*_*x*_, *B*_*y*_, *B*_*z*_, respectively; however, their winding patterns were simpler in comparison. This is because the *B*_*x*_, *B*_*y*_, *B*_*z*_ coils needed to generate a sharp change in the field from a uniform value to zero at the edge of the target region while the *G*_*xz*_, *G*_*yz*_, *G*_*zz*_ coils simply generated a gradually varying field. The six pairs of coils installed in our one-layer magnetically shielded room (Maxshield, Imedco) are shown in **Fig. 2E**.

### 3.2 Theoretical vs Empirical Resistance and Efficiency

The length of the conducting path, the theoretical and measured values of resistance with and without connected segments, and the efficiency of our coils with and without the mu-metal shield are shown in **Table 1**.

**Table 1:** Theoretical and measured properties of the nulling coils. Our geometrically accurate design enables a close match between the theoretical and measured values.

| Coil | Length(m) | | Resistance ($\Omega$) | | | Efficiency (nT/mA or nT/m/mA) | | |
| --- | --- | --- | --- | --- | --- | --- | --- | --- |
|  | Theoretical |  | Theoretical |  | Measured | Theoretical |  | Measured |
|  | bfield<br>-tools | opmcoils | bfield<br>-tools | opmcoils |  | Without<br>shield | With<br>shield |  |
| $B_x$ | 217.4 | 236.5 | 10.7 | 11.6 | 18.2 | 1.5 | 1.4 | 1.4 |
| $B_y$ | 214.3 | 232.2 | 10.5 | 11.4 | 12.8 | 1.4 | 1.3 | 1.3 |
| $B_z$ | 164.5 | 170.6 | 8.1 | 8.4 | 12.5 | 6.4 | 7.1 | 7.1 |
| $G_{xz}$ | 159.3 | 164.9 | 7.8 | 8.1 | 11.8 | 7.5 | 7.8 | 7.9 |
| $G_{yz}$ | 155.8 | 163.2 | 7.7 | 8.0 | 11.8 | 7.8 | 7.9 | 8.4 |
| $G_{zz}$ | 110.1 | 114.6 | 5.4 | 5.6 | 7.7 | 15.4 | 16.4 | 15.4 |

The length of the conducting path in `opmcoils` also considered the length of the connecting segments and their reverse paths and was up to 19 m longer (8% of the total length) than the length of the current loops only. The theoretical resistance agreed with the measured resistance though the measured resistance was systematically higher. The theoretical resistance considering the connecting segments did not fully account for this systematic difference. We hypothesize that the solder joints may be responsible for the discrepancy. However, comparable resistance values has been reported in previous studies (Holmes et al., 2018) which successfully operate OPM nulling coils.

In terms of efficiency, the most efficient coils were the z-direction coils (*B*_*z*_ and *G*_*zz*_) as their optimized design approximates a Helmholtz design. The *B*_*x*_ and *B*_*y*_ coils had a similar theoretical efficiency (1.4 nT/mA). This was expected as the two coils are approximately rotated versions of each other. We noticed that the measured efficiency of the *B*_*y*_ coil was slightly lower (1.3 nT/mA) than the expected theoretical efficiency (1.4 nT/mA). Our initial computation of the theoretical efficiency did not account for the effect of the mu-metal shield. We expected that the high permeability mu metal could impact the measured efficiency. Indeed, since the nulling coils were placed asymmetrically inside the shielded room (closer to the floor) along the y axis, the distortion was largest in this direction. After considering the effect of shielding, the experimentally measured coil efficiency was within 6% of the theoretically measured coil efficiency for all the coils.

### 3.3 Remnant field mapping in target area

The spatial maps of the residual field in the x, y, and z directions after turning on the *B*_*x*_, *B*_*y*_, and *B*_*z*_ nulling coils, respectively, are shown in **Fig. 6**. By manually adjusting the applied current, we were able to reduce the background field to 2 nT. Based on the figure, we did not observe a spatially consistent gradient field that could be cancelled with our gradient field nulling coils. Therefore, we did not apply gradient field nulling for the remaining experiments. Since we operated the sensors in a lightly shielded one-layer room, the background field drifted in time which could affect our estimate of the applied current. Thus, the final background field nulling could be further improved if closed-loop nulling were to be implemented.

### 3.4 Remnant field mapping in helmet

**Fig. 7** shows the spatial map of the residual field along the sensitive axis of the OPM sensor before and after turning on the uniform field nulling coils respectively. The maximum measured background field on the helmet surface was reduced from 21 nT to 2 nT, indicating that the field nulling system could successfully remove the remnant field regardless of the OPM sensor orientation.

**Figure 7:**
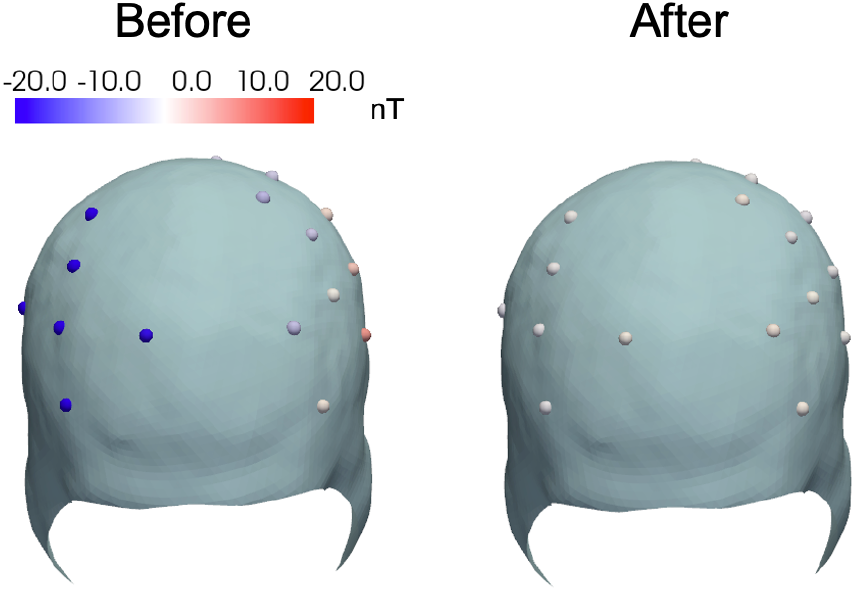
Map of residual magnetic field normal to the helmet surface measured along the sensitive axis of the OPM sensor. The maximum residual field in the OPM sensors after field nulling was 2 nT.

### 3.5 MEG measurement and analysis of somatosensory evoked fields (SEF)

After confirming that we could remove the remnant field on the helmet surface, we used it for a median nerve stimulation experiment using OPM-MEG while the field nulling system was used in the one-layer shielded room. Averaged event-related fields (ERF) in two OPM sensors (N=607) and two SQUID sensors (N=496) from comparable sensor locations are shown in **Fig. 8**. We note that the amplitude of the OPM data was about 2.5 times the SQUID data; however, the noise in the OPM was also proportionally higher. Nonetheless, the OPM measurement was remarkably similar to the SQUID data, showing major ERF components of the somatosensory evoked field (SEF): an early N20 response followed by a P50 deflection and a later component peaking at about 100 ms.

**Figure 8:**
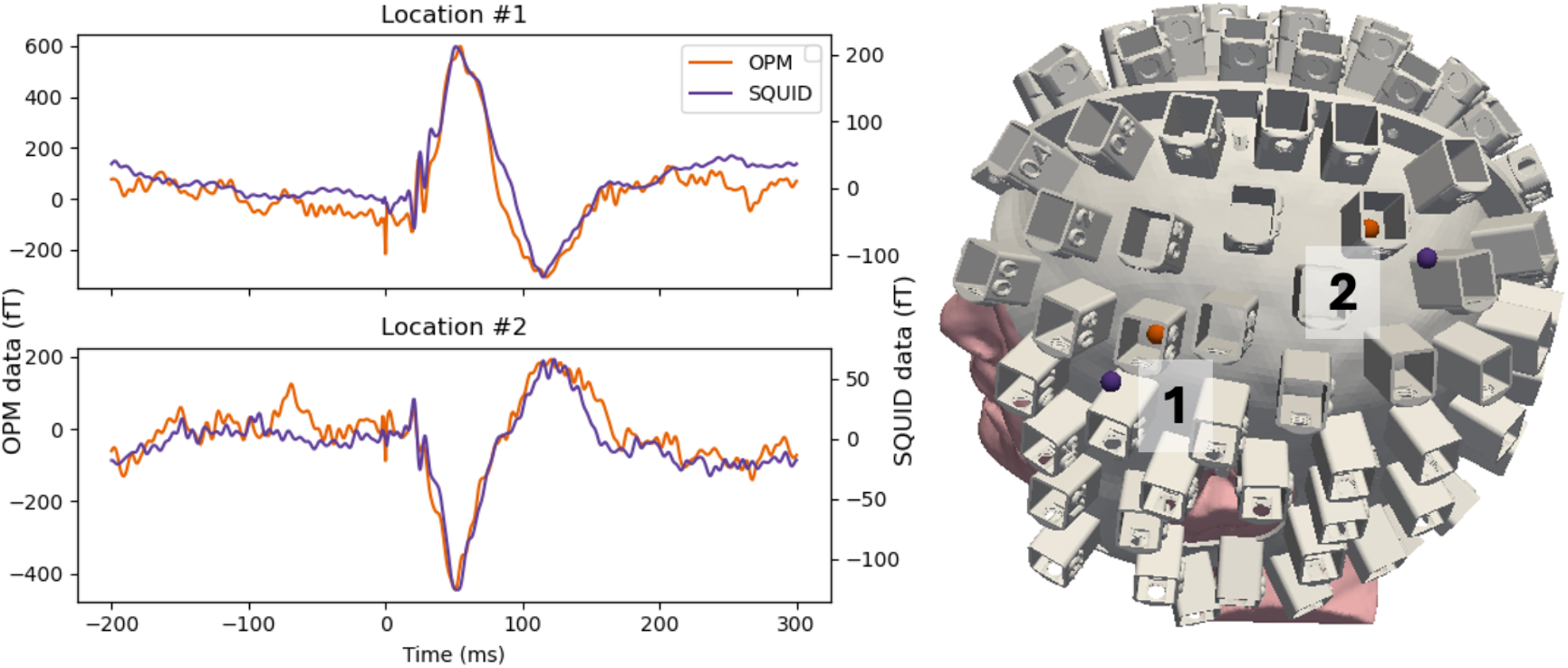
Averaged somatosensory evoked fields (OPM and SQUID) measured at two locations in response to median nerve stimulation (**left**). A subject-specific helmet was created for the experiments by expanding the Freesurfer reconstructed scalp surface (**right**).

### 3.5 Open-source software and hardware

Our work was made possible by the open-source software `bfieldtools` (Mäkinen et al., 2020; Zetter et al., 2020). `Bfieldtools` is a general-purpose software for solving the coil design problem on arbitrary mesh surfaces. Using this software as our base, we implemented additional functionality in `opmcoils` for the practical issues encountered when manufacturing biplanar nulling coils. Our software, `opmcoils` (**Fig. 9**) provides template code to design biplanar nulling coils to null the uniform and gradient components of the background field. The optimized discrete current loops can be connected into a continuous path using an interactive tool and exported to and from KiCad for PCB design. A key feature of our software is that parameters can be provided in physically realizable units: trace width of the PCB in mm, the thickness of the copper layer in oz, and the coil efficiency in nT/mA. Users can also easily compute the impact of the mu-metal shielded room on the coil efficiency (Mäkinen et al., 2020; Zetter et al., 2020). In addition to the software, we also open source the hardware components of our nulling coils. The Gerber files required to print the PCBs, the 3D models used for evaluating them, and the parts list to mount the PCB frame will all be available in our repository. Taken together, we provide users the complete open-source toolkit necessary to download, adjust, and set up PCB-based nulling coils in their MEG centers.

**Figure 9:**
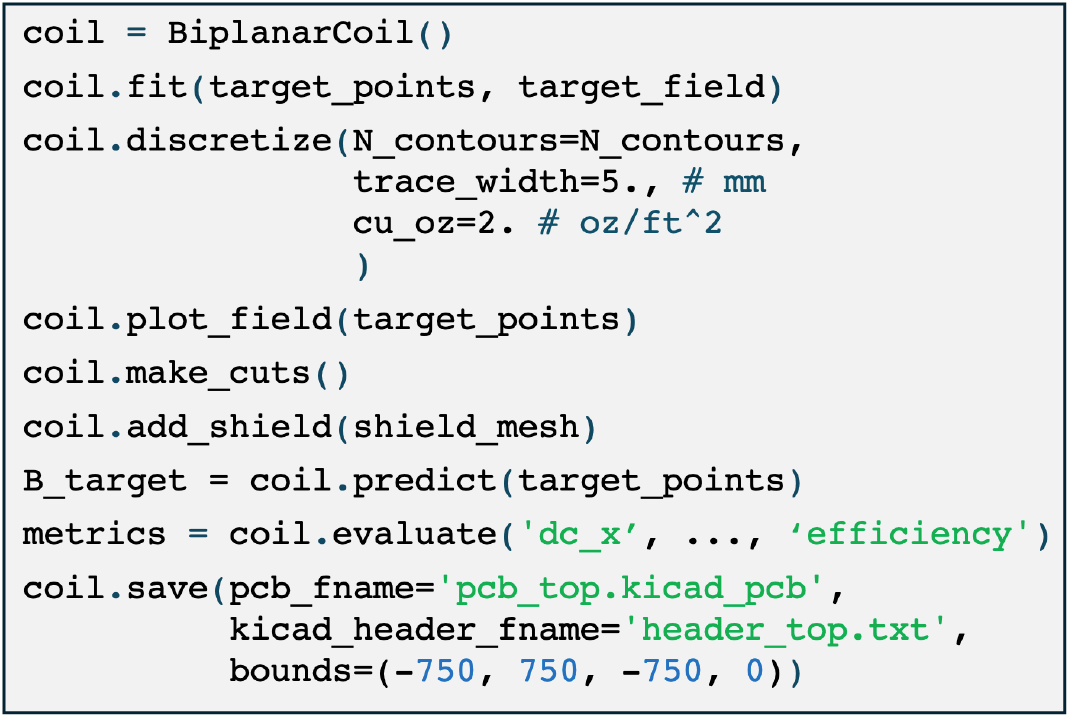
Application Programming Interface (API) for `opmcoils` enabling users to obtain biplanar nulling coils for PCB manufacturing.

## 4. Discussion

### Advantages of open-source PCB design

Manually wound nulling coils have been reported and successfully used in the OPM-MEG literature; however, they are labor intensive due to various challenges in winding the wires. The wire lengths can be over 1000 m and the discrete current loops must be connected into a single continuous path without introducing stray fields. Along with the fact that the winding directions can be either clockwise or anticlockwise, this means that the physical realization of the coils is a complex process. Therefore, it is difficult to be precise, errors are likely, and not easily fixed because the wires are glued on a printed sheet of paper. Another method to construct the coils is to manually wind the coils in 3D printed grooves (McDaniel, 2020). While this technique can be beneficial when the stream function is constrained to a curved surface like an MRI coil, the spatial dimensions for the biplanar coil make 3D printing challenging. It also involves error-prone manual winding.

When we started the design of the nulling coils, easily manufacturable coil designs were not available. Furthermore, the technical implementation details to convert mathematical equations into physical realizations of coils were not easily available. We have streamlined the development process, and we share not only the code used in development but also the coil designs as Gerber files. These Gerber files can be sent to a PCB manufacturer to be printed and used to null fields in any OPM-MEG center. Unlike manually wound nulling coils, PCBs are geometrically precise and easily replicable. Our work can be used to standardize new nulling coil systems for OPM-MEG systems in different environments and test their performance. Ultimately, we hope this will lower the barrier to entry in using OPM-MEG.

Another advantage of our open-source software-hardware PCB design is that the manufacture cost is lower. The open-source approach enables manufacturing the coils at scale and reduces cost for both vendors and users of OPM-MEG systems. Each of the coil pairs costs around $2,000 and the current drivers cost $2,400 each. Including the cost of installation, we estimate that our system cost less than $20,000 in total. We estimate that this is less than one-fifth of the price of commercially available nulling coil systems. By further optimization of the designs, such as a balanced biplanar coil, the cost can be reduced even more (Holmes et al., 2019).

### Design flexibility

In a PCB-based manufacturing system, the coils are chemically etched, the total amount of copper used remains the same, and therefore the cost remains constant regardless of the coil design. Due to this flexibility in design, PCB manufacturing process can be used to design more ambitious coil designs which target higher efficiency or multiple target areas. Indeed, the efficiency can be increased easily by increasing the number of discretized current loops and the resistance can be minimized by increasing the trace width. Even though a high efficiency was not necessary in our shielded room, the design can be readily modified for use in other recording environments with challenging background fields.

### System installation

During installation, care must be taken to route the input wires so that stray fields are minimal. The shielded room can also distort the field and change the efficiency of the coils. The lower edge of our *B*_*y*_ coil was 10 cm from the mu metal floor of our shielded room which reduced the efficiency by 0.1 nT/mA. Raising the coil may improve the efficiency, however the height of the coils was determined by practical considerations such as the height of the subject chair. One can redesign the coil accounting for the effect of the shielded room (Mäkinen et al., 2020; Zetter et al., 2020) but in our case, the measured efficiency was sufficient to operate the OPM-MEG system.

## 5. Conclusions

This work is a demonstration of using PCB-based field nulling systems for OPM-MEG. We successfully designed, developed, and tested the performance of our open-source PCB-based nulling coils for removing uniform background fields and selected spatial gradient components. The PCB design allowed us to obtain empirical efficiency matching the theoretical efficiency to within 6%. The remnant field mapping demonstrated that the background field can be reduced to less than 2 nT after using our nulling coils. We also conducted a median-nerve experiment and demonstrated that the signals obtained using SQUID-MEG and OPM-MEG were comparable when operated in conjunction with our nulling coils. Our open-source field nulling system could lower the barrier to entry for use of OPM-MEG in clinical and neuroscience applications. By leveraging the precision and replicability of PCBs, nulling coil systems can be standardized and evaluated in different environments. In summary, our work streamlines the development process, enables efficient manufacture, reduces costs and manual labor, and improves reproducibility.

## 6. Data and code availability

The software and the Gerber files associated with the nulling coils can be found here: https://opm-martinos.github.io/nulling_coils

## 6. Author contributions

**Mainak Jas:** Conceptualization, methodology, software, validation, investigation, writing - original draft, writing - review and editing, visualization **John Kamataris:** validation, investigation, writing - review and editing **Teppei Matsubara:** visualization, writing - review and editing, investigation, software **Chunling Dong:** methodology, writing - review and editing, investigation **Gabriel Motta:** software, visualization, writing - review and editing **Abbas Sohrabpour:** validation, writing - review and editing, investigation **Seppo Ahlfors:** validation, writing - review and editing, investigation **Matti Hämäläinen:** Conceptualization, writing - review and editing, visualization, funding, investigation **Yoshio Okada** Conceptualization, methodology, validation, investigation, writing - review and editing, funding **Padmavathi Sundaram:** Conceptualization, methodology, software, validation, investigation, writing - review and editing, visualization, funding

## 7. Funding

This work was supported by NIH grants P41EB030006, 1R21NS140619-01, 2R01NS104585-05, 1R01NS112183-01A1, and S10OD030469. The content is solely the responsibility of the authors and does not necessarily represent the official views of the National Institutes of Health.

## 8. Declaration of competing interests

The authors declare that they have no competing interests.

## 9. Acknowledgements

We thank Rasmus Zetter and Lauri Parkkonen for their invaluable insights on the appropriate use of `bfieldtools`, and Jason Stockmann for practical advice on PCB manufacturing.

## Notes

https://opm-martinos.github.io/nulling_coils/

### Competing Interest Statement

The authors have declared no competing interest.

